# Sex differences in osteoblast matrix maturation regulate osteoblast-endothelial interactions

**DOI:** 10.64898/2026.03.11.711110

**Authors:** Aikta Sharma, Roger J.H. Emery, Andrew A Pitsillides, Claire E. Clarkin

## Abstract

**Background:** Bone formation during development and repair is divergently modulated by osteoblast (OB)-derived vascular endothelial growth factor (VEGF) which drives the skeletal sexual dimorphism of the bone vasculature. While the extracellular matrix (ECM) provides both structural and instructive cues to developing vasculature, the contributions of the bone matrix to this skeletal vascular dimorphism in bone remains undefined at the cellular level.

**Methods:** Primary OBs were isolated from neonatal female and male C57BL/6 long bones and cultured under basal or osteogenic conditions. ECM composition was quantified by Raman spectroscopy. Primary murine bone marrow endothelial cells (BMECs) were seeded directly onto established OB layers and maintained in heterotypic cocultures to assess contact-mediated effects of OB ECM on BMEC survival and expansion. OB-conditioned media (CM) was used to evaluate soluble-factor contributions, with VEGF-A concentration quantified by ELISA.

**Results:** Raman spectroscopy, on individual OBs from monotypic cultures, revealed sexually dimorphic ECM signatures that were independent of cellular growth profiles. Female OB matrices were enriched with type I collagen-specific proline and hydroxyproline and octacalcium phosphate with enhanced collagen intra-strand stability consistent with a matrix-dominant signature. Male OB matrices exhibited relatively lower type I collagen content and higher carbonated apatite resulting in an elevated mineral-to-matrix ratio indicative of advanced mineral maturation. After 24-hours of heterotypic culture of BMECs with OBs, BMEC numbers were 1.39-fold higher when in contact with male OBs. CM treatment of BMECs did not recapitulate these effects despite higher VEGF-A release from male OBs.

**Conclusions:** Sex differences in OB ECM are linked to divergent, contact-dependent regulation of BMEC behaviour. These findings suggest that differences in matrix maturation stat contribute to the sex-specific regulation of the skeletal vascular niche. Elucidating the mechanisms that regulate sex-specific OB-ECM production may reveal new therapeutic targets for selectively modulating pathological skeletal angiogenesis in men and women.

**Summary:** Bone is a sexually dimorphic organ, with men and women differing in bone size, strength and risk of fracture. The skeletal vasculature is essential for bone growth and repair, with bone forming osteoblast (OB) cells influencing blood vessel development through the skeletal extracellular matrix (ECM). Although the interactions between OB and vascular cells are crucial for lifelong skeletal health, it is not yet known whether sex differences in bone structure between men and women arise from differences in OB activity, or whether this divergence is driven by sex differences in blood vessel growth. Here, we show that male and female mouse OBs deposit distinct ECMs that differentially influence vascular endothelial cell behaviour. Female OBs produce a collagen-rich matrix with low mineral content. In contrast, male OB matrices contain less collagen and more mineral while releasing elevated levels of blood vessel promoting VEGF-A than females. When placed directly onto these OBs, vascular cell growth was greater when in contact with male than female OBs. Notably, this sex-dependent effect requires direct contact between both cell types and was not reproduced by exposure to OB-derived substances alone. These findings identify a cellular mechanism by which sex differences in OB matrix composition influences vascular cell behaviour in bone. Understanding how OB-vascular interactions differ by sex may help explain variation in bone health, healing capacity and disease risk between men and women. Further, our approach may support the discovery of new therapeutic targets that support bone growth and repair while targeting abnormal blood vessel growth in a sex-specific manner.

**Highlights:**

- Primary OBs from male and female C57BL/6 mouse long bones synthesise compositionally distinct ECMs.
- Female OB matrices are type I collagen-rich and enriched with octacalcium phosphate, whereas male OB matrices contain less type I collagen and higher levels of carbonated apatite.
- Bone marrow-derived endothelial cell (BMEC) growth is enhanced in heterotypic cocultures with male, but not female, OBs after 24 hours.
- Male OBs release higher levels of the pro-angiogenic factor VEGF-A than female OBs.
- The sex-specific effects of the OB ECM on BMECs is contact-dependent and are not reproduced by treatment with OB-derived conditioned media.

## Introduction

The skeletal system is profoundly sexually dimorphic, with differences in bone structure, strength and geometry emerging during puberty and becoming amplified with age under the influence of gonadal sex hormones [1–4]. During skeletal growth, males undergo greater periosteal apposition, resulting in greater bone diameter and cortical thickness. In contrast, during puberty, females experience relative inhibition of periosteal apposition and instead increase cortical thickness through endosteal bone formation thereby reducing the endosteal perimeter. These sex-specific growth trajectories contribute to larger bone size, increased bone strength and lower fracture rate in men than women [1–3, 5]. However, variations in body and bone size alone do not fully account for sexually dimorphic skeletal phenotypes [6] and instead suggest that additional regulatory mechanisms contribute to these differences.

Circulating sex steroids are central regulators of bone homeostasis [7, 8], yet their cellular effects are not uniform between the sexes due to the involvement of divergent receptor-mediated pathways within skeletal cell populations. Estrogen receptor α (ERα) is a principal mediator of estradiol-stimulated bone formation [9–13] in both sexes [14] whereas androgen receptor [15–17] signalling is critical for male skeletal accrual and maintenance [16]. Increasing evidence indicates that sexual dimorphism in bone is not solely hormone driven [18, 19]. Indeed, intrinsic sex differences have been reported across a number of skeletal cell types including OBs [20], osteoclasts [21, 22] and osteocytes [23, 24], implying that cell-autonomous mechanisms that may be influenced by sex chromosomes, epigenetic programming or early developmental imprinting contribute to the lifelong skeletal divergence. For instance, osteoclast precursor populations exhibit sex differences, with female cells undergoing accelerated osteoclastic differentiation alongside transcriptional enrichment of genes involved in innate immune and inflammatory response pathways [21]. OBs derived from human craniosynostosis patients similarly display sex-specific differences in differentiation, with male OBs showing higher early differentiation activity and reduced proliferation compared with females [25].

Bone vascularisation is essential for mineralisation, nutrient delivery and osteogenic coupling [26] and is partly regulated by OB-derived vascular endothelial growth factor (VEGF) [27, 28]. Despite advances enabling detailed visualisation of skeletal vascular networks [29–32], whether sex differences in bone matrix composition contribute to the regulation of skeletal endothelial cell function remains poorly defined. We previously showed that OB-specific deletion of *Vegfa* (VEGFKO) produces sexually dimorphic vascular and matrix phenotypes in vivo [33, 34], and that *Vegf*-deficient OBs synthesise sex-specific extracellular matrices (ECMs) in vitro with impaired mineralisation in males associated with increased vascularisation in vivo [33]. Given the tight coupling between osteogenesis and angiogenesis [31, 35–37], we hypothesised that intrinsic sex differences in OB matrix assembly directly modulate skeletal endothelial cell behaviour. To test this, we employed separate heterotypic primary coculture models comprising male and female OBs with bone marrow derived endothelial cells to determine whether sex-specific matrix composition directly modulates vascular endothelial cell behaviours. By integrating functional coculture assays with Raman spectroscopic profiling of matrix composition, we investigated whether intrinsic sexual dimorphism in OB behaviours is sufficient to drive sex-dependent endothelial responses to thereby contribute to the sexually dimorphic regulation of the skeletal vascular niche.

### Sex and gender terminology

In accordance with sex and gender reporting guidelines (SAGER), the terms *woman/women* and *man/men* used in this paper to refer to individuals identified as biologically male or female. For rodent experiments, *female* and *male* refer exclusively to biological sex.

## Results

### Sex differences in pro-angiogenic VEGF release and matrix deposition in OBs are independent of OB growth profiles

Primary OBs were isolated from the long bones of 4-5 day-old male and female C57BL/6 mouse pups and cultured for 7 days under basal or osteogenic (osteo) conditions prior to the characterisation of sex-specific ECM composition by Raman spectroscopy (Figure 1A). As demonstrated previously, Raman spectroscopy enables the detection of matrix and mineral components of the ECM synthesised by differentiating primary OBs, corresponding with elevated alkaline phosphatase activity and positive Alizarin Red staining [38]. At day 7 of culture, no overt sex differences in OB morphology were observed under either culture condition (Figure 1B), with OBs derived from both sexes consistently adopting a flattened appearance with elongated processes. Growth curve analysis confirmed similar proliferation profiles between male and female cultures across timepoints and media conditions, with cell number increasing in both populations over time as expected (Figure 1C and D, respectively). Despite comparable growth, quantification of VEGF-A released into CM by ELISA revealed a 1.26-fold increase in VEGF-A levels from male OBs cultured under osteo conditions compared to female OBs (P<0.0001) (Figure 1D). No significant sex differences in VEGF-A release were detected under basal conditions.

**Figure 1:**
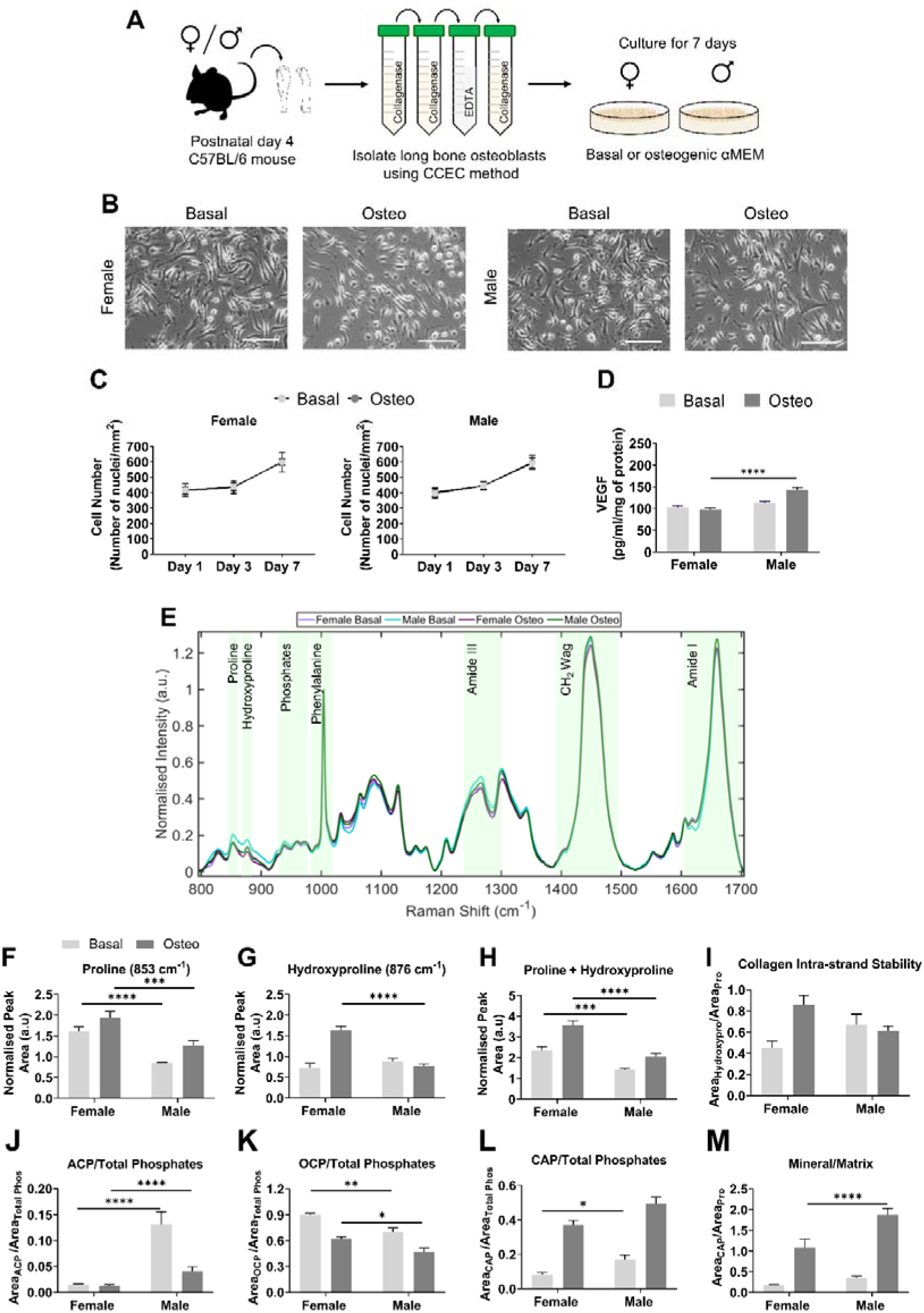
Sex differences in pro-angiogenic matrix deposition in OB cultures are independent of cellular growth profiles. Primary long bone osteoblasts (OBs) were isolated from 4-day old male and female C57BL/6 mice using the collagenase-collagenase-EDTA-collagenase (CCEC) method ahead of monolayer subculture in tissue culture plastic or quartz coverslips for 7 days under basal or osteogenic (osteo) culture conditions (A). Representative phase contrast images show similarities between male and female OB phenotype when cultured in basal and osteo conditions at day 7 (B, scale bars represent 100 µm). Cell number increased in time in culture however no significant differences in the growth profiles of female (B) versus male OBs (C) were detected on day 1, 3 and day 7 in between culture conditions. OB-derived VEGF release was elevated in male versus female OBs cultured under osteo conditions (D). Spectrum-wide deviations in class means of normalised Raman spectra (N=125 spectra per group) were observed in bands corresponding to ECM (proline, 853 cm^-1^; hydroxyproline, 876 cm^-1^; amide III region, 1242 cm^-1^; CH_2_ wag, 1450 cm^-1^; amide I, 1660 cm^-1^) and mineral components (phosphate region, 940 cm^-1^ to 980 cm^-1^) between female and male OB cultures (A). Spectral deconvolution of Raman spectra highlighted sex-differences in type I collagen components (proline, B; hydroxyproline, C; and their sum, D), in female OBs but no sex differences were evident in the intra-strand stability of the collagen triple helix (E). Further divergence was noted in the mineral precursors of hydroxyapatite, amorphous calcium phosphate (F, ACP), octacalcium phosphate (G, OCP) and carbonated apatite (H, CAP) between the sexes and are reflected in the mineral/matrix ratio, describing overall matrix maturity (I). Data presented as mean normalised value ± SEM from three individual biological replicates. Interaction between sex and culture condition was assessed using a two-way ANOVA with further statistical significance between groups evaluated using Tukey’s post-hoc test (*P<0.05, **P<0.01, ***P<0.001, ****P<0.0001).

### Raman spectroscopy reveals sex-specific matrix and mineral signatures of male and female OBs

Raman spectroscopy was performed on day 7 OB cultures to characterise ECM composition. Spectral measurements were acquired from OB populations cultured under basal and osteo conditions within the “Fingerprint region” ranging from 800 cm^-1^ to 1700 cm^-1^ (Figure 1E). Inspection of class mean spectra within this region revealed clear sex differences in the intensity of Raman peaks and bands corresponding to type I collagen components including proline (853 cm^-1^) and hydroxyproline (876 cm^-1^), CH_2_ deformation (1450 cm^-1^) and amide I (1660 cm^-1^), in addition to mineral-related phosphate vibrations spanning 940 cm^-1^ to 980 cm^-1^.

Univariate spectral deconvolution, performed on normalised class mean spectra, revealed significantly higher proline peak area in female OB versus male OB cultures under both basal (P<0.0001) and osteo (P=0.0006) conditions (Figure 1H). The peak area of hydroxyproline was likewise elevated in female OBs, with statistical significance only reached in OBs cultured under osteo conditions (P<0.0001, Figure 1G). The combined peak area of proline and hydroxyproline (853 cm^-1^ + 876 cm^-1^, Figure 1H), used as a surrogate for total collagen content, was also consistently higher in female than male OB cultures irrespective of media type (basal, P=0.0001 and osteo, P<0.0001). In contrast, the hydroxyproline/proline ratio, determined by the area ratio of the hydroxyproline to proline peak and used to assess the collagen intra-strand stability, did not differ between the sexes (Figure 2I). To assess whether these observed sex differences were also evident in mineral-associated Raman bands, further spectral analysis was conducted into the mineral precursors of hydroxyapatite as a function of the total phosphate region. Amorphous calcium phosphate (ACP, 948 cm^-1^), the earliest detectable precursor of hydroxyapatite, was significantly higher in male than female OB cultures under both basal and osteo conditions (both P<0.0001, Figure 1J). Conversely, the intermediate octacalcium phosphate (OCP, 970 cm^-1^) phase was enriched in female OBs cultures across both media conditions (basal, P=0.002 and osteo, P=0.03; Figure 1K). Carbonated apatite (CAP, 959 cm^-1^), representing the final mineral precursor of hydroxyapatite, was elevated in basal male OB cultures (P=0.05), with a similar trend towards elevated CAP levels also evident in osteo culture conditions (Figure 1L). Consistent with these findings, the mineral/matrix ratio (CAP/proline peak area) was significantly greater in osteo male OB cultures compared to females (P<0.0001, Figure 2M). Together, these spectral analyses reveal that female OBs predominantly deposit a collagen-dominant, OCP-enriched matrix whereas male OBs generate a comparatively mineral-enriched ECM characterised by increased abundance of ACP and CAP and an elevated mineral/matrix ratio reminiscent of a more advanced stage of mineral maturation.

**Figure 2:**
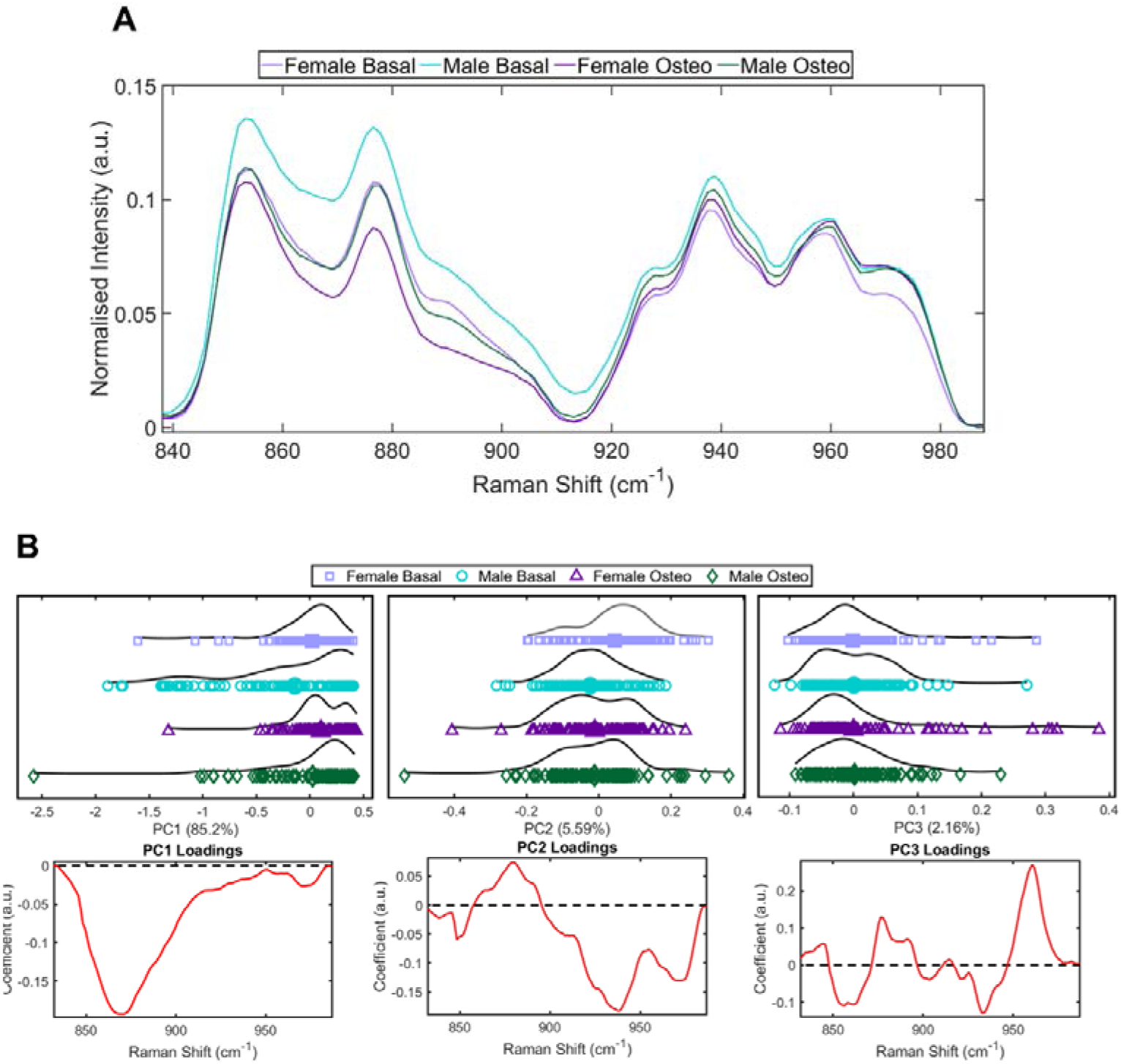
PCA analysis of Raman spectra enables distinction of ECM components between the sexes and culture conditions. Zoomed class means Raman spectra of spectral regions corresponding to ECM components from 840cm^-1^ to 980 cm^-1^ (A) (N=125 spectra per condition). PCA analysis, presented as 1-Dimensional scatter plots, show distribution of Raman peaks and bands that account for the greatest variation (PC1>PC2>PC3) for female OBs cultured in basal (lilac, square) and osteo conditions (purple, triangle) and male OBs cultured in basal (blue, circle) and osteo conditions (green diamond) for 7 days (B). Mean loading score is shown as a single larger symbol for each condition, with smaller symbols representing the PC score for a single spectrum. Corresponding PCA loading plots representing the coefficient of variation beneath represent spectral regions that contribute the highest variation.

Osteogenic induction exerted further sex-dependent effects on ECM composition (Table 1). For type I collagen-specific species, osteogenic stimulation resulted in increased proline peak area in male OB cultures whereas no corresponding change was observed in female cultures. In contrast, increases in the peak area of hydroxyproline were only evident in osteo female OB cultures. Although the sum peak area of hydroxyproline and proline was significantly increased in response to osteo induction in both female and male OBs, the magnitude of this change was greater in female cultures. Consistent with this, the collagen intra-strand stability ratio was also significantly higher in female OBs cultured under osteo conditions compared with basal controls, while remaining comparable across male OB cultures. The abundance of mineral precursor species was also differentially influenced by osteo culture. ACP peak area decreased exclusively in osteo male OB cultures, while the peak area of OCP decreased in both sexes compared to basal controls. Two-Way ANOVA confirmed significant sex x media interactions for the hydroxyproline peak area, collagen intra-strand stability ratio, ACP peak area and the mineral/matrix ratio, whereas no interaction was detected for the proline, summed proline and hydroxyproline peak area, OCP or CAP peak area. Collectively, these data indicate that female OBs exhibit a collagen-dominant and OCP-enriched signature whereas male OBs display a comparatively mineral-enriched ECM profile characterised by higher ACP and CAP abundance and an elevated mineral/matrix ratio.

**Table 1:** Effect of culture condition on ECM composition in female and male OB cultures. Table summarising the effect of culture in basal and osteo conditions on the spectral composition of ECM components synthesised by female and male OB culture. Interaction between sex and culture condition was assessed using a two-way ANOVA with further statistical significance between groups (basal versus osteo) evaluated using Tukey’s post-hoc test (*P<0.05, **P<0.01, ***P<0.001, ****P<0.0001).

|  | Female Basal<br>vs<br>Female Osteo |  | Male Basal<br>vs<br>Male Osteo |  | Sex and<br>Media<br>Interaction |
| --- | --- | --- | --- | --- | --- |
| Peak assignment | P value | Effect | P value | Effect | P value |
| Proline | NS | ↔ | * (P=0.05) | ↑ | NS |
| Hydroxyproline | **** (P<0.0001) | ↑↑ | NS | ↔ | ****<br>(P<0.0001) |
| Proline + Hydroxyproline | **** (P<0.0001) | ↑↑ | * (P=0.03) | ↑ | NS |
| Collagen Intra-strand Stability Ratio | ** (P=0.002) | ↑ | NS | ↔ | **<br>(P=0.004) |
| Amorphous Calcium Phosphate | NS | ↔ | **** (P<0.0001) | ↓↓↓ | **<br>(P=0.002) |
| Octacalcium Phosphate | **** (P<0.0001) | ↓↓ | *** (P=0.0003) | ↓↓ | NS |
| Carbonated Apatite | **** (P<0.0001) | ↑↑↑ | **** (P<0.0001) | ↑↑↑ | NS |
| Mineral/Matrix Ratio | **** (P<0.0001) | ↑↑↑ | **** (P<0.0001) | ↑↑↑ | * (P=0.03) |

In order to further interrogate sex-dependent spectral variation, principal component analysis (PCA) was applied to the 840 to 980 cm^-1^ regions of the class means spectra (Figure 2A). PC1 accounted for 85.2% of the total variance and clearly differentiated Raman spectra according to sex and media condition, with separation primarily driven by the proline-associated peak at ∼870 cm^-1^ (C-C stretching mode; Figure 2B, left). Here, male basal OB spectra (mean score = -0.14163) clustered distinctly from female basal (PC mean score = 0.0175935), female osteo (PC mean score = 0.101326) and male osteo (PC mean score = 0.0253243) OB spectra, consistent with biochemical divergence in ECM composition. PC2 (5.59% variance; Figure 2B, middle) was predominantly influenced by the by the ∼879 cm^-1^ hydroxyproline peak, distinguishing female basal spectra (PC mean score = 0.0433638) from male basal (PC mean score = -0.0233738), female osteo (PC mean score = -0.0107576) and male osteo (PC mean score = -0.0105736) OB spectra with minor contributions from peaks at ∼848 cm^-1^, ∼937 cm^-1^ and ∼971 cm^-1^ corresponding to polysaccharide, protein and C-C wagging modes, respectively. PC3 accounted for 2.16% of the variance (Figure 2B, right) and provided minor additional separation. Female basal (PC mean score = -0.0014436) and female osteo (PC mean score = -0.000250587) spectra clustered together, driven by contributions from ∼856 cm^-1^ (proline) and 932 cm^-1^ peak (C-C bond vibration), whereas male basal (PC mean score = 0.000317781) and male osteo (PC mean score = 0.00159337) OB spectra were distinguished by peaks at ∼877 cm^-1^, ∼892 cm^-1^ and ∼960 cm^-1^ corresponding to lipids, phosphodiester and CAP, respectively.

### Contact-mediated survival of BMECs is divergently influenced by OB sex in coculture

To determine whether the OB-derived ECM influences endothelial cell behaviour in a sex-specific manner, fluorescently labelled primary murine BMECs were seeded onto established male and female OB monolayers on day 7 and maintained in direct-contact coculture (50:50 OB:BMEC medium) for up to 72 hours (Figure 3A; timeline in Figure S1). BMEC survival was assessed using fluorescence microscopy.

**Figure 3:**
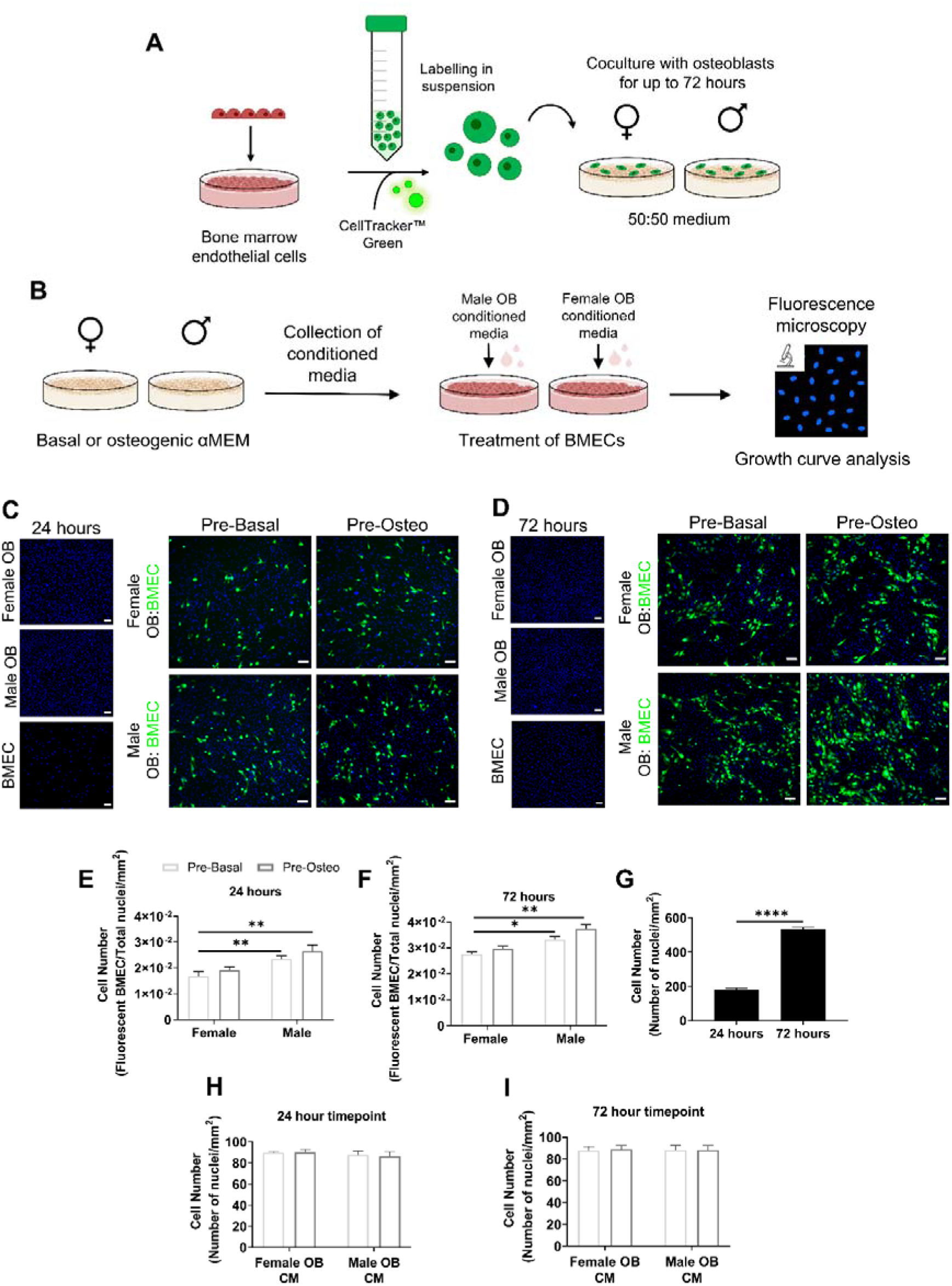
The survival of BMECs is supported when cocultured with male OBs. Primary mouse bone marrow endothelial cells (BMECs) were labelled in suspension using CellTracker Green™ before being added to male and female OB cells and maintained in direct-contact cocultures (1:1) in 50:50 OB:BMEC media for up to 72 hours (A) in preparation for fluorescence microscopy-based analyses. Parallel female and male OB monocultures cultured in basal and osteo conditions for were established for conditioned media collection on day 7 of culture and used to treat BMECs for up to 72 hours ahead of growth curve analysis (B). Representative fluorescence images of nuclei stained with Hoechst 33258 in female OB, male OB and BMEC monolayers on day 8 (24 hour timepoint, C) and day 10 (72 hour timepoint, D) are shown alongside the corresponding cocultures with CellTracker™ Green labelled BMECs for each timepoint and OB culture condition (scale bars represent 50 µm). The number of BMECs when cocultured with male OBs, presented as a function of the total cell number of nuclei in each coculture, were elevated following 24 hours of coculture on day 8 (E), with further elevations evidenced after 72 hours (F). Quantification of the growth profiles of BMECs show increase in cell number over time (G). Treatment of BMEC monocultures with conditioned media, collected from female and male OB monocultures on day 8 (H) and day 10 (I), had no effect on BMEC number. Data presented as mean normalised cell number ± SEM from three individual biological replicates. Statistical significance between conditions was assessed using two-way ANOVA and Tukey’s post-hoc test for interactions between sex and culture condition (E, F, H and I) or student’s t-test (G) or (*P<0.05, **P<0.01, ***P<0.001, ****P<0.0001).

Following 24 hours of direct-contact heterotypic coculture, sex differences in BMEC number were evident (Figure 3C), and became more pronounced by 72 hours (Figure 3D). Quantification of cell numbers after 24 hours of coculture revealed that BMEC number increased when cocultured with female and male OBs preconditioned by osteo media for 7 days (both P=0.005, Figure 3E). By 72 hours, BMEC growth was more strongly supported in coculture with male OBs under both basal (P=0.04) and osteo conditions (P=0.007) compared to females (Figure 3F). To assess whether these differences were matrix-dependent, BMECs were maintained in monoculture in 50:50 medium for 72 hours. As expected, BMEC number increased over time (P<0.0001), reaching levels comparable to coculture conditions (Figure 3G). However, treatment with BMEC monocultures with CM collected from male and female OB monocultures at equivalent 24-hour and 72-hour coculture timepoints (Figure 3H and I) did not alter BMEC number. Likewise, exposure to 50:50 OB:BMEC media did not affect the growth profiles of monocultured female or male OBs (Figure S1B and C, respectively). Collectively, these findings indicate that enhanced BMEC expansion in coculture is contact-dependent and occurs in a sex-specific manner, whereas soluble factors alone are insufficient to recapitulate these effects.

## Discussion

Sexual dimorphism in bone geometry, strength and fracture susceptibility is well established, yet the cellular mechanisms that link OB function to sex-specific vascular regulation remain incompletely understood. In our previous work, OB-derived VEGF was shown to divergently modulate mineralisation in males and females leading to sexually dimorphic vascular phenotypes at the tibiofibular junction. In the present study, we demonstrate that male and female primary OBs deposit compositionally distinct ECMs which divergently support the survival of vascular endothelial cells and thereby contribute to sexually dimorphic angiogenic microenvironments in bone.

Using heterotypic coculture models, we demonstrate that OB sex drives sex-specific responses in endothelial cell in a contact-dependent manner. Direct-contact cocultures revealed enhanced BMEC survival when cocultured with male OBs compared to female OBs under both basal and osteogenic conditions over 72 hours (Figure 4). Intriguingly, this divergence was not reproduced following treatment with OB-conditioned media, despite increased VEGF-A release from male OBs under osteogenic stimulation. This indicates that soluble factors alone are insufficient to account for the observed vascular differences and instead implicate the OB-derived ECM as a critical regulator of endothelial cell behaviour. The deposition of bone matrix and its subsequent mineralisation by OBs occurs as a temporally regulated process and are critical determinants of bone mechanical competence [39, 40]. At the material level, the organic matrix constitutes one third of its mass and two thirds of its volume, providing flexibility and resistance to tension and torsion [41]. At the compositional level, type I collagen consists of triple-helical polyproline II-type chains comprised of repeating Xaa-Yaa-Gly motifs wherein Xaa and Yaa are often proline and hydroxyproline (28% and 35%, respectively) [42, 43]. The mineral phase, conversely, dictates the elastic modulus, yield stress and ultimate stress of bone [44, 45] and forms the remaining two thirds of the mass of bone and third of its volume [41]. Mineral, comprising calcium and phosphate ions are accommodated by the collagenous ECM during matrix mineralisation, and is recognised to take a number of precursor forms before its progressive maturation into carbonated apatite [46–49].

**Figure 4:**
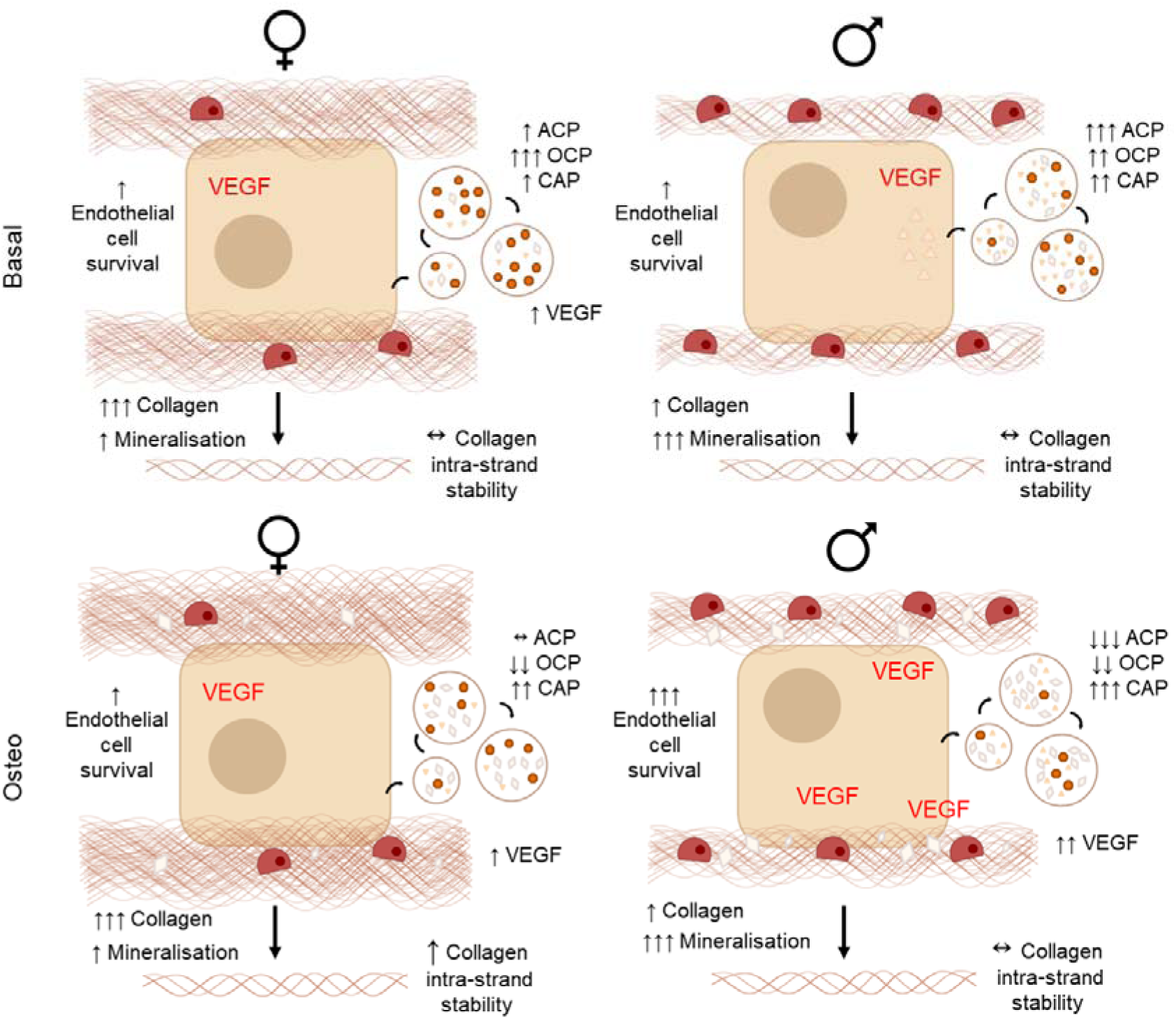
Sexual dimorphism in the matrix composition of OBs drives divergence in BMEC survival. In female OB cultures (left), Raman spectroscopy revealed that the ECM was dominated by collagen-specific species, with high intra-strand stability coupled to low levels of mineralisation while male OB cultures (right) were characterised by low levels of collagen-specific species and comparatively weaker intra-strand stability with high levels of mineralisation in osteo culture (bottom). Pro-angiogenic VEGF release was elevated in male OB cultures, which alongside the ECM composition appeared to support BMEC survival in direct contact cocultures relatively versus that observed in female OB:BMEC cocultures. These sex-differences are indicative of a direct sex-specific role of matrix composition in the regulation of BMEC activity and survival.

In order to define the compositional basis of our observed OB-ECM-dependent effects, we employed Raman spectroscopy as this approach enables simultaneous assessment of organic and inorganic components of the bone matrix [33, 34] and has been shown to correlate well with bending stiffness compared to conventional measures of bone mineral density (BMD) [50]. Our analyses revealed marked sexual divergence in ECM composition, with female OB-ECMs consisting of increased type I collagen-specific proline and hydroxyproline accompanied by the mineral precursors ACP and OCP. This contrasted with male OB matrices which comprised lower levels of these collagen-associated species yet were enriched with CAP. The predominance of OCP in female OB cultures coupled with increased type I collagen is indicative of active osteoid deposition transitioning into the early stages of matrix mineralisation. Within the same time frame and independently of cell number, male OBs appear to progress towards later mineralisation stages despite reduced collagen deposition compared to female OBs. These disparities in ECM signatures indicate that matrix deposition and maturation kinetics differ between the sexes while the structural quality of collagen remains comparable. This interpretation is supported by the comparable hydroxyproline/proline ratios observed between the sexes, indicating that post-translational modifications of collagen are preserved despite differences in matrix abundance. These compositional differences align with established sex differences in skeletal growth trajectories, wherein males undergo prolonged cortical bone accrual through periosteal expansion while females experience earlier skeletal maturation with preferential endosteal bone formation [1, 51, 52]. In mice, this is further substantiated by reports of reduced expression of *Alpl* and *Bglap*, ALP activity and nodule formation in female OBs cultures derived from C57BL/6 mice alongside reduced differentiation potential of bone marrow stromal cells from female C57BL/6 mice compared with males [53]. Critically, the presence of these sexually divergent matrix signatures in vitro, in the absence of circulating sex hormones supports the concept that OBs retain this autonomous sex-specific functional programming ex vivo, which has similarly been reported in other bone cell populations including osteoclasts [21, 22] and osteocytes [23, 24].

However, beyond functional implications, these sex differences in matrix composition appear to contribute to divergent vascular responses within the skeletal niche. Despite possessing a collagen-rich matrix, female OB cultures exerted a comparatively weaker pro-survival influence on BMECs than male OBs cultures. This observation suggests that collagen abundance alone is insufficient to support endothelial survival and instead implicates additional matrix-associated factors including mineral-phase composition and VEGF-ECM interactions. Consistent with this interpretation, male OB cultures exhibited greater VEGF release under osteogenic culture conditions. Given that specific VEGF isoforms bind to ECM components [54, 55], and that growth factor delivery in conjunction with CAP enhances angiogenic responses during bone regeneration [56, 57], it is plausible that CAP-enriched matrices produced by male OBs facilitate the local retention or presentation of pro-angiogenic signals at the OB-BMEC interface. Indeed, skeletal angiogenesis is closely coupled to osteogenesis and is reflected by the spatial intimacy maintained between OBs and vascular endothelial cells throughout embryonic development and postnatal growth [58, 59]. OB-derived VEGF functions through paracrine, autocrine and intracrine signalling mechanisms to stimulate endothelial proliferation and vessel formation via VEGF receptor activation [28, 58, 60]. However, endothelial cells also require adhesion to the ECM to support migration, survival and angiogenic organisation. In bone and other tissues, type I collagen has been widely reported to promote angiogenesis both in vitro and in vivo [61–66]. For example, endothelial cells cultured on type I collagen matrices undergo rapid cytoskeletal reorganisation and form branching capillary-like networks [62, 67]. Our findings contrast with this paradigm and instead suggest that endothelial behaviour reflects the integrated biochemical environment of the matrix and may be influenced by the presentation of pro-angiogenic cues from local cell types and their matrix components.

Although intrinsic sex differences in OB matrix composition have not previously been described, sexually dimorphic regulation of the ECM has been reported across a number of biological systems. For example, sex differences in ECM composition and tissue mechanics have been observed in the brain, where neuroinflammatory ECM remodelling has been shown to contribute to altered cortical viscoelasticity in multiple sclerosis [68, 69]. Similarly, sex-specific matrix remodelling has been reported in cardiovascular tissues, where the ageing male cardiac ECM exhibits enrichment of distinct collagen isoforms associated with pro-fibrotic, injury-like remodelling phenotypes [70, 71]. At the cellular level, sexually divergent ECM regulation has also been described in vascular cell populations, including valvular interstitial cells and smooth muscle progenitor cells where differences in collagen synthesis, matrix remodelling enzymes and proliferative behaviours have been described to contribute to sex differences in ECM composition and disease susceptibility [72, 73]. A clearer understanding of the mechanisms governing sex-specific matrix assembly at the cellular level in skeletal tissues is therefore important for elucidating how bone matrix organisation and angiogenic processes are differentially regulated in males and females. Investigating these interactions in vivo remains challenging as the bone matrix is highly mineralised and as a consequence, the mechanisms linking OB-derived matrix properties to vascular behaviour have been difficult to study. The nanoscale differences in matrix composition described herein may explain the sex-specific gender trends in the prevalence of a number of skeletal pathologies including osteopenia [74], osteoarthritis [75], osteoporosis [76] and fracture repair [77] which are increasingly recognised to involve systemic vascular alterations [78, 79]. Defining how OB-vascular crosstalk is modulated by matrix composition may therefore inform more accurate diagnosis and the development of sex-specific therapeutics, improved design of biomaterials or bone regenerative approaches that are tailored to distinct skeletal microenvironments.

A key limitation of the present study is that the molecular mechanisms underlying the sexual divergence in BMEC survival observed in the direct contact, heterotypic co-cultures could not be determined. Although our findings demonstrate a contact-dependent enhancement of endothelial survival in cocultures with male OBs, the downstream signalling pathways responsible for these effects remain to be elucidated. Potential mechanisms may involve differential integrin-mediated adhesion, VEGF receptor signalling or matrix stiffness-dependent mechanotransduction arising from distinct ECM compositions identified here. Future studies could address these questions by isolating these OB and BMEC populations from cocultures using fluorescence activated cell sorting based on CD31 expression, enabling downstream transcriptomic or targeted gene expression analyses of angiogenesis-related pathways, including matrix remodelling enzyme expression, VEGF receptor signalling and integrin expression. In addition, in vivo studies will be necessary to determine how these intrinsic matrix differences interact with systemic hormonal environments during skeletal development and ageing.

In conclusion, this study demonstrates that OBs from male and female mice deposit intrinsically distinct ECMs that differentially regulate endothelial cell behaviour through contact-dependent mechanisms. Raman spectroscopic analyses revealed that female OBs generate a collagen-rich ECM enriched in early mineral precursors whereas male OBs produce matrices with reduced type I collagen abundance and increased CAP, indicative of a more mineral-mature ECM. These compositional differences were accompanied by enhanced BMEC survival in cocultures with male OBs and was supported by greater OB-derived VEGF release in this sex suggesting that sex-specific matrix assembly contributes to divergent angiogenic microenvironments in bone. By situating these findings within the broader context of sexual dimorphism in skeletal growth and vascular biology, our work identifies OB matrix programming as a cell autonomous contributor of sex-specific regulation of the skeletal vasculature. Together, these findings provide a means to link nanoscale matrix composition to sexually dimorphic skeletal architectures and may inform the development of sex-specific regenerative therapeutic strategies.

## Materials and Methods

### Isolation and culture of male and female OBs

All mice utilised in these studies were kept in controlled conditions at the Biomedical Research Facility, University Hospital Southampton, UK (*ad libitum* access to food and water, controlled temperature (20 °C to 24 °C), humidity (55% ± 10%) and light cycles (12 hours on and 12 hours off)). All cell isolation procedures were performed in accordance with the UK Animals (Scientific Procedures) Act of 1986 and regulations set by the UK Home Office and local institutional guidelines. Primary long bone OBs were separately isolated from 4-5 day old male and female C57BL/6 mouse littermates, from four individual litters using established methods as previously described [80]. Isolated cells were resuspended in basal αMEM containing 10% (v/v) heat-inactivated fetal bovine serum (FBS), 0.1% gentamicin and 100 U/ml penicillin, 100 µg streptomycin for culture in 75 cm^2^ flasks. Cells were cultured until 80% confluent in a humidified incubator set at 37 °C/5% CO_2_ and used at passage 1.

At confluence, male and female OBs were seeded on uncoated quartz coverslips (#No5, thickness: 0.5 mm, Ø: 20 mm; UQG-Optics, UK) at a density of 10,000 cells per well and maintained in monolayers as previously described [38]. In preparation for heterotypic direct-contact coculture, male and female OBs were plated at a density of 10,000 cells per well, with corresponding mono-culture controls seeded at the same time and density. All OBs were seeded in basal αMEM and left for adhere for 48 hours. Following this, further culture was performed in basal, or osteogenic (osteo) media comprised of basal 10% (v/v) heat-inactivated FBS supplemented 50 μg/ml L-ascorbic acid and 2 mM β-glycerophosphate [38, 80]. Culture media were replenished every third day until day 7 of culture unless otherwise stated.

### Protein assay and VEGF ELISA

On day 6 of OB culture, the cell culture media were aspirated and replaced with low serum basal αMEM containing 1% (v/v) FBS. Cells were cultured further for 24 hours ahead of conditioned media (CM) collection on day 7 of culture. The Pierce™ bicinchoninic acid (BCA) protein assay kit and reagents (Thermofisher Scientific, USA) were used to quantify total protein concentration in cell lysates, isolated as previously described [81].

A VEGF-A mouse sandwich enzyme linked immunosorbent assay (ELISA) kit and reagents (R&D Systems, USA) was used to quantify natural and recombinant VEGF (VEGF_120_ and VEGF_164_) in the collected CM as per the manufacturers protocols. VEGF concentrations quantified in the conditioned media were normalised to total protein concentration.

### Raman spectroscopy, spectral processing and analysis

Following CM collection on day 7, male and female OBs cell layers were briefly washed with 1X PBS prior to fixation in 4% formaldehyde in preparation for Raman spectroscopy. Raman spectra were collected using the InVia® Raman microscope (Renishaw, UK) using the 532 nm laser (with a Gaussian beam profile and 2400 lines/mm diffraction grating, providing spectral resolution of ∼1.06 cm^-1^) and 63X water immersion objective (Leica, Germany) with a numerical aperture of 1.2 (providing a diffraction limited spot size of 280 nm) as previously described [38]. Wavenumber and intensity calibration were performed to the 520.7 cm^-1^ peak of an internal silicon wafer ahead of spectral acquisition. Raman spectra were acquired using single point static scans with an exposure time of 20 seconds, three accumulations and 100% laser power within the 800 cm^-1^ to 1700 cm^-1^ range of the “fingerprint region” [82] in conjunction with the Renishaw® WiRE 4.1 software and a charged coupled device. Raman spectra were collected from 5 points within the cytoplasmic region of 25 male and female OBs per group across three biological replicates (corresponding to individual OB isolations, Figure S2A-D). Raman spectra of blank quartz were also acquired using the same spectral acquisition settings for post-acquisition background subtraction procedures. Power at the sample was approximately 30 mW.

Cosmic ray artefacts were removed from individual spectra upon acquisition and quartz contributions removed using manual background subtraction procedures as previously described [38]. Class means spectral denosing (Haar-wavelets, 6-point smoothing), background correction (9^th^-order polynomial fitting) in addition to wavenumber and intensity normalisation (1004 cm^-1^) were performed in iRootLab [83, 84] for spectral comparisons between sexes and culture conditions. Raman peaks and bands of interest, including those corresponding to the ECM (proline, 853 cm^-1^ and hydroxyproline, 876 cm^-1^), and the mineral phase (amorphous calcium phosphate, ACP, 948 cm^-1^; octacalcium phosphate, OCP, 970 cm^-1^ and carbonated apatite, CAP, 959 cm^-1^) were identified in second order derivative spectra (Figure S2E) in preparation for peak area extraction through univariate spectral deconvolution in WiRE 4.1 (Figure S2F) as previously described [38]. For multivariate analysis, spectra were pre-processed as described above then mean centred for principal component analysis (PCA) with a maximum of 10 principal components in iRootLab [83, 84]

### Culture of bone marrow endothelial cells

Primary BMECs (Generon, UK) were cultured in complete basal endothelial cell media and utilised at passage 3 to 5 following the manufacturer’s protocols and recommendations.

### Fluorescent labelling of bone marrow endothelial cells

CellTracker™ Green 5-chloromethlfluorescein diacetate dye, prepared in dimethylsulfoxide (DMSO) (Invitrogen, UK) was utilised for the fluorescent labelling of BMECs following the manufacturer’s instructions. 1×10^6^ BMECs were treated with the CellTracker™ (5 µM in serum-free endothelial cell media) in suspension for 30 minutes at 37 °C. Control cells were prepared following the same procedure in serum-free basal endothelial cell media containing equivalent volumes of DMSO. Further cell dilutions were made in complete basal endothelial cell media containing ahead of plating.

### Direct contact heterotypic coculture

After 7 days of OB culture, the culture media were aspirated and replaced with fresh basal αMEM containing 10% FBS. 10,000 unlabelled and CellTracker™ Green-labelled BMECs were then plated directly into same wells containing the male and female OBs, in complete endothelial cell media equivalating to 50:50 OB:BMEC media per well (Figure 3A). Heterotypic cell layers were maintained in direct contact cocultures for 24 and 72 hours in a humidified incubator set at 37 °C/5% CO_2_. 72 hours was selected as the endpoint for this series of experiments due to the substantial decay of CellTracker™ fluorescence (Figure S3).

Coculture control wells consisted of monocultures of OBs alongside unlabelled and CellTracker™ Green-labelled BMECs at density of 10,000 cells per well. BMEC monocultures were seeded in complete endothelial cell media and left to adhere for 2 hours. Following this, equivalent volumes of basal αMEM was added to give 50:50 OB:BMEC media per well as described above. Experimental nomenclature used here forth is as follows: pre-basal corresponds to OBs that were cultured in basal medium for 7 days prior to BMEC addition, pre-osteo corresponds to the OBs that were cultured in osteogenic medium for 7 days prior to BMEC addition (Figure 1 and Figure S1A).

### Quantification of BMEC survival

After 24 hours and 72 hours of coculture, the culture media were aspirated, and cell layers were briefly washed with 1X PBS prior to fixation with 4% formaldehyde. Cell nuclei were fluorescently labelled with Hoechst 33258 (Invitrogen, UK) at a concentration of 1 µg/ml for 15 minutes in the dark at room temperature. Cocultured cells were imaged using the Deltavision Elite fluorescence microscope (GE Healthcare Sciences, USA) in combination with the SoftWoRx software (version 6). Dual-channel images of Hoechst fluorescence and the CellTracker™ Green fluorescence were captured sequentially using the in-built DAPI bandpass filter set (excitation: 390 ± 18 nm and emission: 435 ± 48 nm) and the bandpass filter set for fluorescein isothiocyanate (FITC) detection (excitation: 475 ± 28 nm and emission: 525 ± 48 nm), respectively, using a 10X air objective with numerical aperture of 0.4. The imaged field of view was 1.33 mm x 1.33 mm. The number of CellTracker™ labelled BMECs, containing a visibly blue-fluorescent nucleus, were manually counted using the “multi-point” function in FIJI [85] after background subtraction, intensity normalisation and pseudocolouring (BMECs, green; nuclei, blue). The total number of cell nuclei in each image were determined using an automated custom in-house macro in FIJI as previously described [33]. The number of CellTracker™ Green labelled BMECs were subtracted from the number of nuclei in each individual pair of DAPI/FITC fluorescence microscopy images originating from the same field of view to determine the number of OBs in the overlaid images. The number of fluorescent BMECs were then divided by the number of OBs to give the number of BMECs per OB prior to normalisation to the imaged field of view. A total of 40 images were acquired randomly from four individual cocultures per condition from each biological replicate.

### Growth curves

To assess the growth profiles of OBs over time across the culture conditions, cell layers were fixed in 4% formaldehyde on day 1, 3, 7, 8 and 10 of culture. To monitor the growth of BMECs, the same procedure was followed however cell layers were fixed after 24 and 72 hours of culture in line with the coculture time points. Following fixation, all cell layers were washed with 1X PBS and cell nuclei were stained with Hoechst 33258 and subsequently imaged and analysed as described above.

### Conditioned media collection and treatment

CM were collected from each OB monoculture at the corresponding 24 hour and 72 hour coculture time points and used to treat unlabelled BMECs, plated at density of 10,000 cells per well in basal endothelial cell media for 24 hours. CM-treated BMECs were subsequently fixed with 4% formaldehyde, stained with Hoechst 33258 and fluorescently imaged prior to growth curve analysis as described above).

### Statistical analysis

All data are presented as mean value ± SEM. The effects of culture condition in male and female OB and BMEC datasets was statistically evaluated using one-way analysis of variance (ANOVA) or paired student’s t-test where appropriate. Interaction between sex and culture condition was assessed using a two-way ANOVA with Tukey’s post hoc test for further comparisons. All statistical analysis was performed using the GraphPad Prism (version 9) (GraphPad Software, USA). P values of less than 0.05 were considered to be statistically significant and are noted as *. P values of <0.01, <0.001 and <0.0001 are denoted as **, *** and **** respectively.

## Notes

### Competing Interest Statement

The authors have declared no competing interest.

