## Supplementary Information for "Sex differences in osteoblast matrix maturation regulate osteoblast-endothelial interactions"

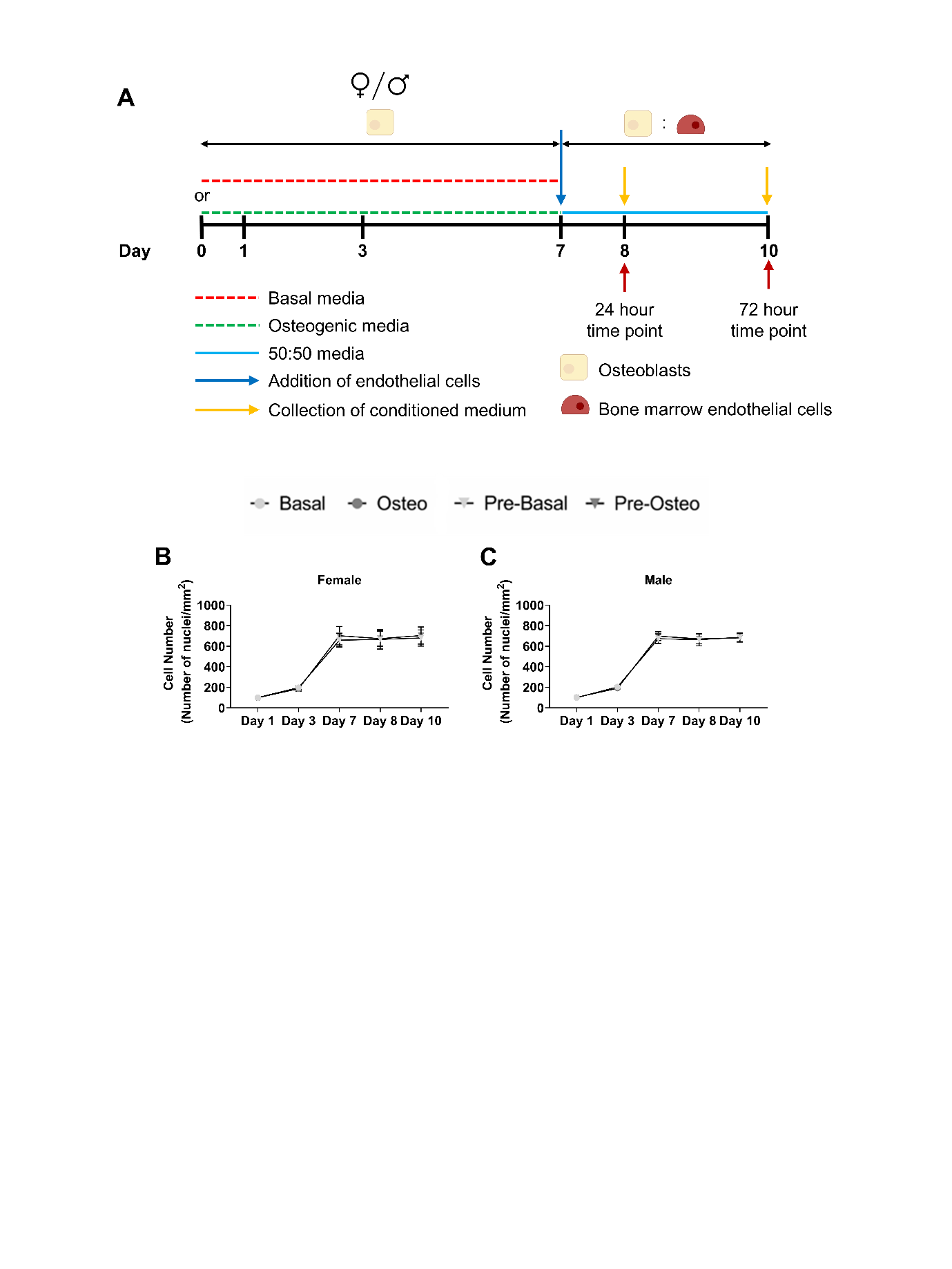

**Figure S1:** Experimental timeline for coculture studies (A). Female and male primary long bone OBs, extracted from 4-day old C57BL/6 mice were cultured under basal (red dashed line) or osteo (green dashed line) for 7 days, On day 7, the culture media was removed and BMECs were introduced in 50:50 OB:BMEC media, at a 1:1 seeding ratio. Direct contact cocultures were further maintained for 24 hours (day 8), and 72 hours (day 10) ahead of quantitative analysis. Cellular growth profiles of female and male OBs were comparable in basal and osteo media (day 1 and 3) and in 50:50 OB:BMEC media (day 7, 8 and 10) with time (B and C).

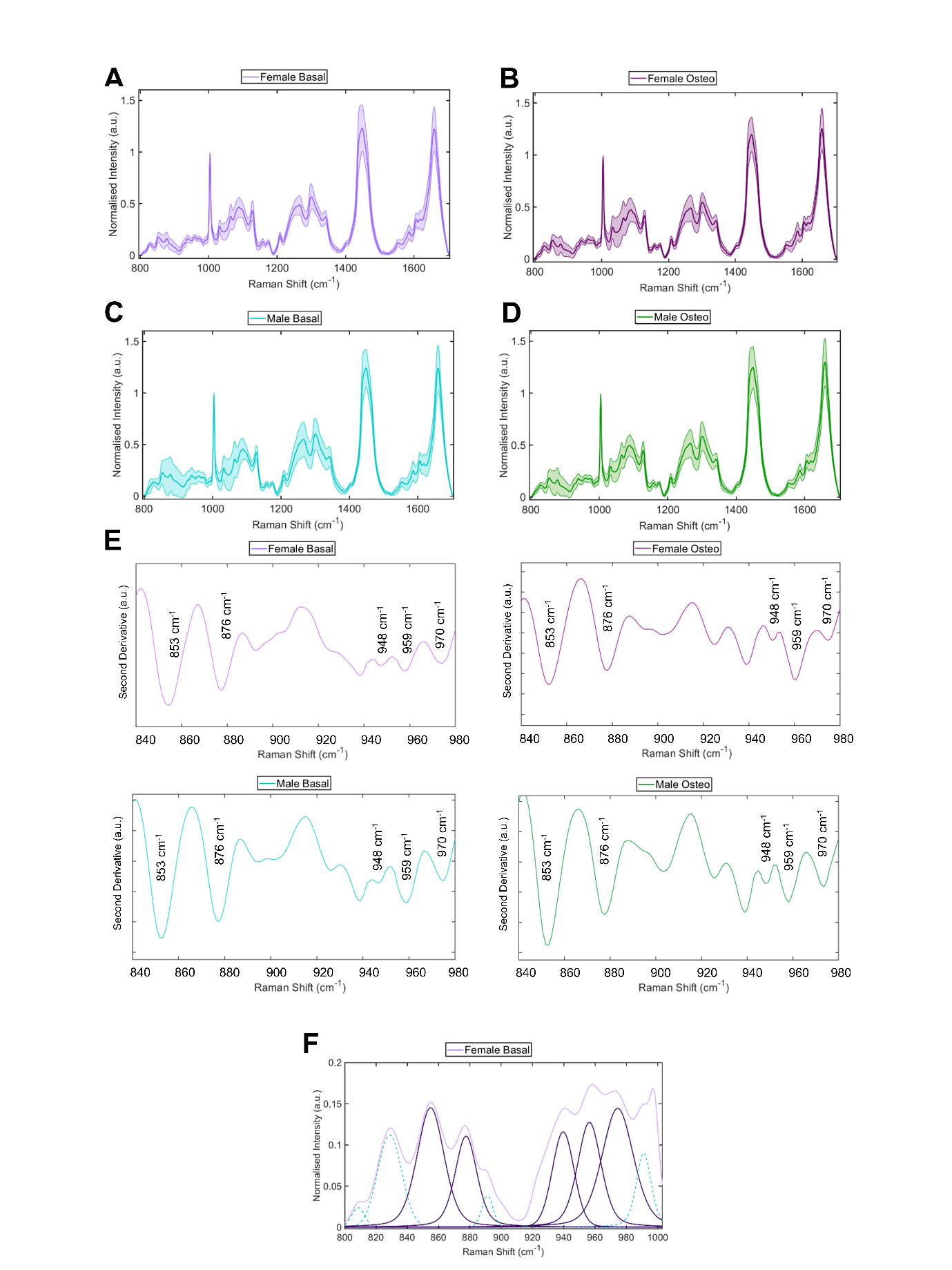

**Figure S2:** Raman spectra collected within the fingerprint region from 800 cm^-1^ to 1700 cm^-1^, presented as class means of n=125 single spectra with standard deviation displayed, obtained from female (A, B) and male OBs (C, D). Second derivative spectra, indicating position of bands of interest including those arising from the ECM (proline, 853 cm^-1^ and hydroxyproline, 876 cm^-1^) and mineral species within the phosphate region (940 cm^-1^ to 980 cm^-1^) (E), were obtained from female and male OBs for each culture condition and used for spectral deconvolution analyses. Exemplary peak fitting of class mean spectra following second derivative calculation are shown (F), with curves corresponding to peaks of interest shown as opaque lines.

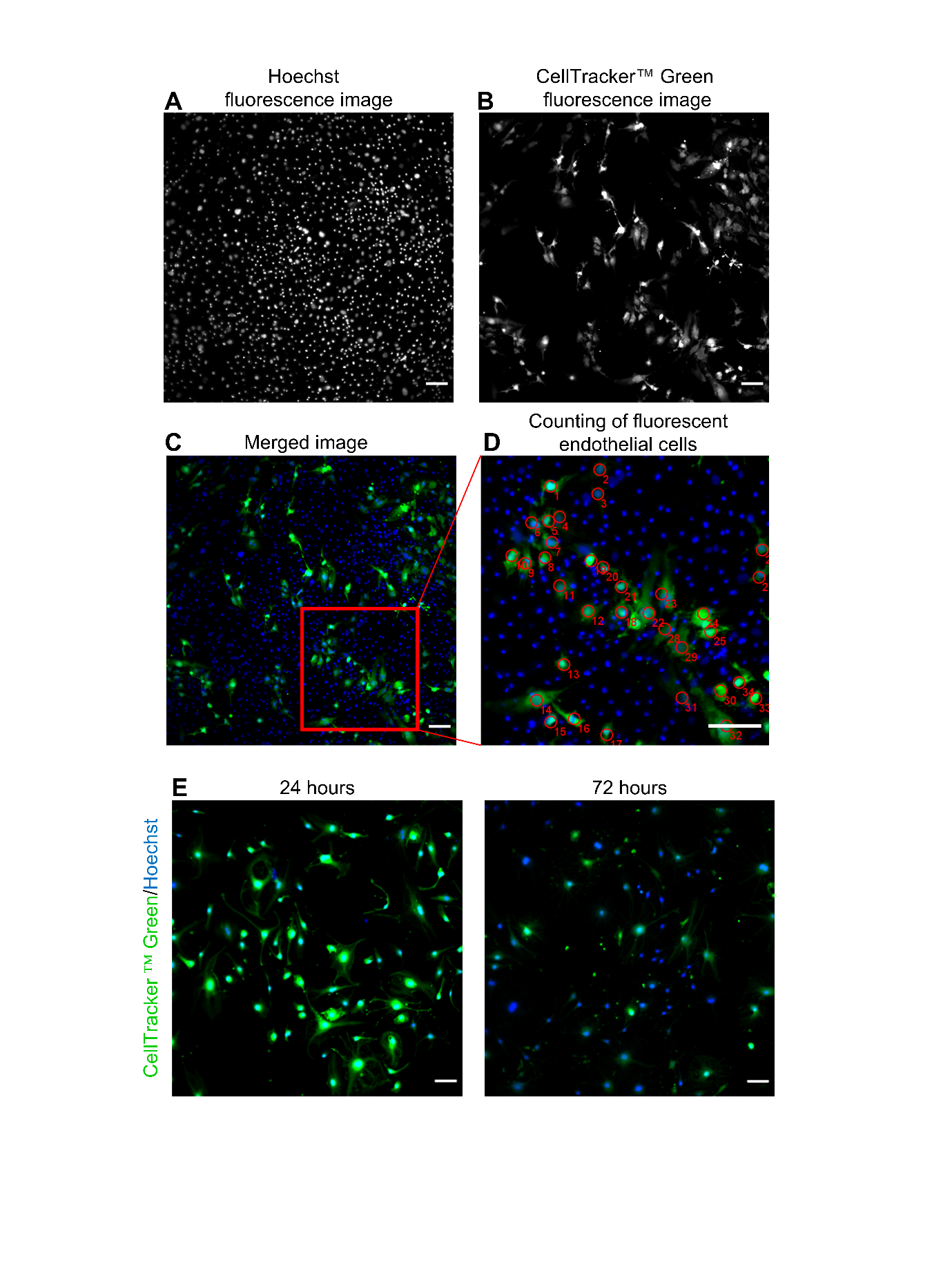

**Figure S3:** Visualisation and calculation of the number of CellTracker™ Green labelled BMECs for cell survival analysis. Fluorescence microscopy of cocultures containing CellTracker™ Green labelled BMECs (A) and Hoechst 33258 labelled nuclei (B) were obtained sequentially, from the same field of view, pseudocoloured and overlaid (C; nuclei, blue and BMEC, green). The number of fluorescently labelled BMECs were quantified by manual counting of green-fluorescent cells containing a visible blue-fluorescent nucleus in Fiji (D). Decay in CellTracker™ Green fluorescence was apparent following 24 hours of culture (E). Scale bars represent 50 µm.
